# A Literature-Based Knowledge Graph Embedding Method for Identifying Drug Repurposing Opportunities in Rare Diseases

**DOI:** 10.1101/727925

**Authors:** Daniel N. Sosa, Alexander Derry, Margaret Guo, Eric Wei, Connor Brinton, Russ B. Altman

## Abstract

One in ten people are affected by rare diseases, and three out of ten children with rare diseases will not live past age five. However, the small market size of individual rare diseases, combined with the time and capital requirements of pharmaceutical R&D, have hindered the development of new drugs for these cases. A promising alternative is drug repurposing, whereby existing FDA-approved drugs might be used to treat diseases different from their original indications. In order to generate drug repurposing hypotheses in a systematic and comprehensive fashion, it is essential to integrate information from across the literature of pharmacology, genetics, and pathology. To this end, we leverage a newly developed knowledge graph, the Global Network of Biomedical Relationships (GNBR). GNBR is a large, heterogeneous knowledge graph comprising drug, disease, and gene (or protein) entities linked by a small set of semantic themes derived from the abstracts of biomedical literature. We apply a knowledge graph embedding method that explicitly models the uncertainty associated with literature-derived relationships and uses link prediction to generate drug repurposing hypotheses. This approach achieves high performance on a gold-standard test set of known drug indications (AUROC = 0.89) and is capable of generating novel repurposing hypotheses, which we independently validate using external literature sources and protein interaction networks. Finally, we demonstrate the ability of our model to produce explanations of its predictions.

## 1. Introduction

In the United States, rare diseases are defined as diseases that affect fewer than 200,000 people each. Although individually rare, the cumulative effect of all rare diseases amounts to a significant proportion of the population there are an estimated 7,000 rare diseases that affect 25–30 million Americans.^1^ A major challenge of rare disease research is that despite this aggregate health burden, no single rare disease affects enough people to be prioritized for drug development over other, more prevalent diseases. As a result, there has historically been a lack of academic and pharmaceutical research for rare disease treatments, and the vast majority of rare diseases still have no therapeutic options. One way to address this unmet clinical need is through drug repurposing, or the use of pharmaceuticals already existing in the market to treat different diseases than they were developed to treat. This paradigm has been successful in many contexts with examples including methotrexate and sildenafil (Viagra).^2,3^ Previously, drug repurposing has largely been accomplished by clinical observation of drug side effects, but a systematic, data-driven approach for identifying repurposing opportunities is needed to improve efficiency and coverage.

Advancements in computation and machine learning have enabled natural language processing (NLP) techniques that are effective and scalable for processing large bodies of unstructured text. Recently, NLP was applied to all *∼*28.6 million PubMed abstracts to synthesize and summarize the relationships between drugs, genes/proteins, and diseases into a heterogeneous knowledge graph known as the Global Network of Biomedical Relationships (GNBR).^4^ This dataset is powerful because (1) it is large, consisting of over 130,000 entities and over two million edges, (2) each of these edges is represented by a set of several important, semantic themes, and (3) the confidence associated with each of these themes is quantified as a continuous value. By harnessing GNBR, we can synthesize disparate sources of knowledge relevant to rare diseases in order to systematically generate repurposing hypotheses that can be directly mapped back to the literature.

Previous data-driven approaches to drug repurposing have relied on gene expression, chemical structure, or electronic health records data.^5^ For example, a gene expression–based drug repurposing method was described by Sirota et al. to repurpose topiramate as a therapeutic option for inflammatory bowel disease.^6^ Traditional network-based methods have focused on identifying disease modules and using diffusion strategies to rank novel interactions.^7,8^ Recently, network embedding methods, which learn a mapping from nodes and edges to low-dimensional vectors such that the proximity structure of the original network is preserved in the embedding space, have attracted great interest. The resulting vectors provide an ideal platform for machine learning tasks, and have been applied in pharmacological applications such as the prediction of polypharmacy side effects and drug-drug interactions.^9,10^ Building on such successes, we implement a network embedding method for drug repurposing that explicitly models the confidence of relationships in GNBR based on their evidence in literature.^11^ This model is more appropriate for representing and learning from literature-based knowledge, which is inherently noisy. As far as we know, we are the first to incorporate such uncertainty into a literature-based graph embedding method, allowing for a more precise and nuanced drug repurposing model. Unlike previous methods, our hypotheses do not rely on any curated databases, allowing the model to automatically improve as the volume of literature proliferates and GNBR expands.

In this work, we first prioritize rare diseases based on their potential for drug repurposing, accounting for the availability of data and the current state of treatment need. Then, we develop a knowledge graph embedding–based drug repurposing method that produces treatment hypotheses with strong evidence in literature and evaluate our results using gold-standard drug indications. We then apply our model to generate novel drug repurposing hypotheses and assess their scientific validity using a variety of sources. Finally, for top-scoring hypotheses we elucidate recurring network patterns that contribute to our predictions and demonstrate their capacity to provide mechanistic interpretations.

## 2. Methods

### 2.1. Data

GNBR contains edges or links between two entities from among the set {gene, drug, disease} and a support score (normalized between 0 and 1) representing the literature-derived confidence of the relationships between those two entities.^4^ The relationships are divided into 32 high-level semantic themes (Fig. 1) and are organized into four categories based on the entities they connect. For example, the edge between metformin and type 2 diabetes, a well-established relationship, has a support for the theme “Treatment” of 0.999. Some themes exist in multiple categories. In some cases, the two entities have a single very clear relationship, and in others, there is literature evidence for several relationships.

**Fig. 1:**
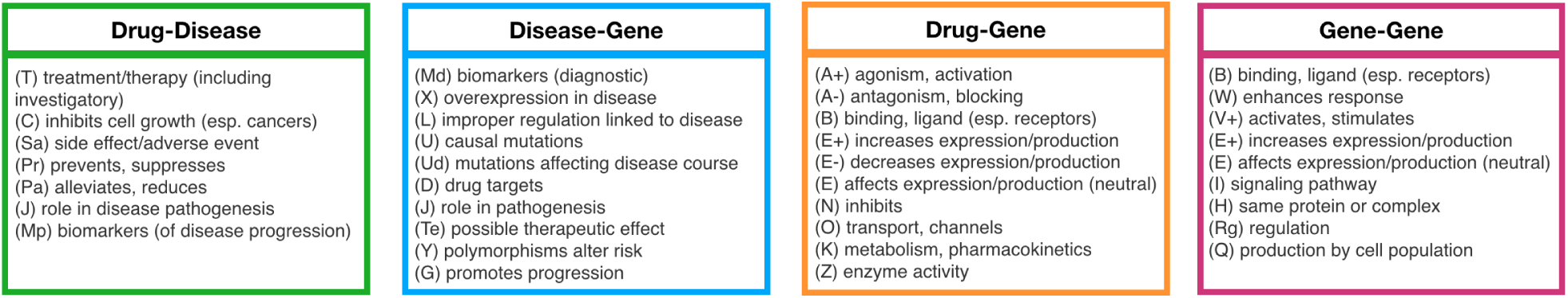
Summary of all themes in GNBR, organized by category along with their reference codes.

Using rare disease information from Orphanet,^12^ we extracted MeSH,^13^ OMIM^14^ and UMLS^15^ IDs for each rare disease in Orphanet. By directly matching MeSH and OMIM IDs and indirectly matching UMLS IDs using the UMLS Metathesaurus, we identified 2,779 rare diseases in GNBR. We maximize clinical utility of our method by focusing on diseases with high prevalence and no FDA-approved indications. Prevalence was retrieved from Orphanet, and FDA-approved indications are found on DrugCentral.^16^ Finally, we filtered GNBR by identifying the largest connected component of the graph after removing any node that is not (1) a gene node, (2) a high-priority rare disease node, or (3) a drug/disease node present in a known indication. The resulting graph comprised a total of 63,252 nodes and 583,685 edges.

### 2.2. Embedding-based prediction method

We adopt an uncertain knowledge graph embedding method,^11^ which takes advantage of the support scores in GNBR (i.e. the confidence of the relationship) in order to learn embedding vectors for all nodes and themes. The geometric intuition for this model is that the proximity between the vectors for head *h*, relation *r*, and tail *t* is related to the confidence score associated with the triple (*h, r, t*). Concretely, for a given triple *l* = (*h, r, t*), a plausibility score, *g*, is defined by the corresponding embeddings ***h, r, t*** as follows:

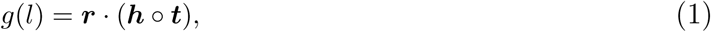

where ○ denotes the element-wise product. This score is mapped to the interval [0, 1] through the bounded rectifier function

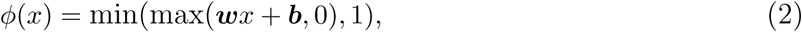

where ***w*** and ***b*** are learned parameters. The final predicted confidence score, *f* (*l*), is thus:

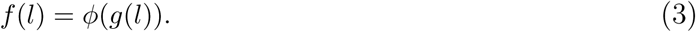

In order to accurately rank candidate triples and avoid ties, we remove the min-max bounding in Eq. (2) at test time. For every triple used in training, a corresponding “negative” triple is sampled by corrupting the tail node and resampling a random node with an assumed support of 0. The joint objective function to be minimized is the sum of squared errors between the prediction, *f* (*l*), and support, *s*_*l*_, for each triple *l*:

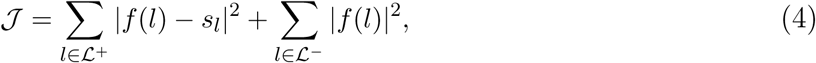

where *ℒ*^+^ is the set of triples in GNBR and *ℒ*^−^is the set of sampled negative triples. We formalize the generation of drug repurposing hypotheses as a link prediction task in which we predict high-confidence triples of the form (*Drug*, “Treatment”, *Disease*) using the learned embeddings.

## 3. Experiments and Results

### 3.1. Embedding-based predictions

#### 3.1.1. Experimental Design

As an internal validation, we quantify the ability for our model to recapitulate known gold standard drug-disease indications in the embedding-based link prediction task. For this we use MEDI,^17^ a database of drug indications compiled from SIDER 2,^18^ RxNorm,^19^ MedlinePlus,^20^ and Wikipedia. Drug-disease combinations were mapped from ICD9 codes to UMLS codes, resulting in 3,329 combinations comprising 811 drugs and 360 diseases.

Triples in GNBR containing a known indication pair from MEDI were split into 60% training, 20% validation, and 20% test sets. All other triples in GNBR were split into 90% training, 5% validation, and 5% test sets. We chose an embedding dimensionality of *d* = 128, and trained the model for 100 epochs with batch size 1024. We used the Adam optimizer for training, with exponential decay rates *β*_1_ = 0.9 and *β*_2_ = 0.99.^21^ The validation set was used to determine early stopping criteria based on mean squared error and to tune the learning rate *lr* ∈ {0.001, 0.005, 0.01}.

To ascertain which parts of the network were most valuable to the embeddings, we considered three submodels: (1) removing all drug-disease triples with relationships other than “Treatment”, (2) removing all gene-associated triples and taking the largest connected component from the resulting drug-disease bipartite graph, and (3) considering only triples of the form (*Drug*_*i*_, “Treatment”, *Disease*_*j*_) without embedding the network at all. Our test set consisted of 355 MEDI drug indications as positives and 355 randomly-sampled pairs as negatives, where the drugs and diseases were drawn from the sets of all drugs and diseases present in the known indications.

#### 3.1.2. Performance on gold-standard indications

To assess the capacity of our model to recapitulate known treatment indications, we calculated the receiver operating characteristic (ROC) and precision-recall (PR) curves for all submodels based on the predicted confidence score given in Eq. (3). Our full embedding model performs well in discriminating between the positive and negative pairs, achieving an area under the ROC curve (AUROC) of 0.89 (Fig. 2). Removing non-Treatment drug-disease themes decreases performance markedly, as expected because the other drug-disease themes are semantically related and often correlated in support scores. Therefore, many training examples that would positively contribute to the final embeddings are lost. The “Treatment” theme alone achieves an AUROC of 0.83, indicating that the support score for that theme is in fact a suitable proxy for confidence that a true indication exists. However, this submodel fails to capture indirect relationships and thus cannot predict new links. Performance increases slightly when the gene-related triples are removed from the network, most likely because the embeddings for drugs and diseases are no longer constrained to be consistent with the genes, which dominate the number of triples in training.

**Fig. 2:**
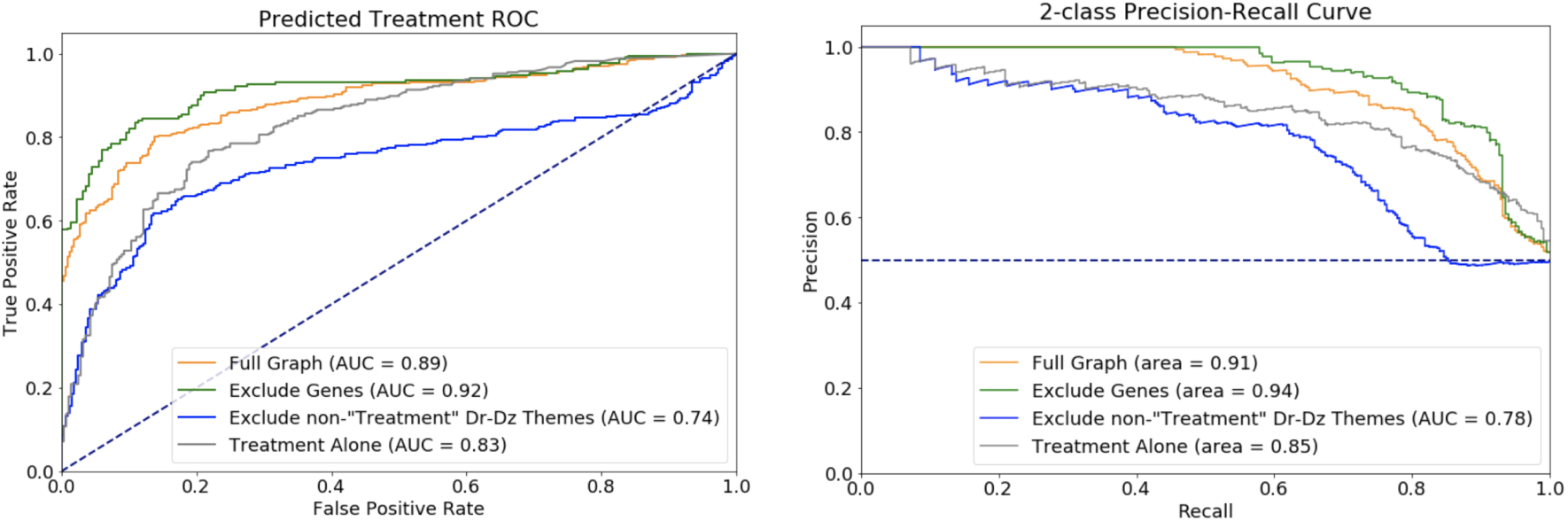
Treatment prediction on gold-standard test set for different submodels, including ROC (left) and PR (right) curves.

However, this is less useful for novel drug repurposing because it takes advantage of only transitive relationships between drugs and diseases and fails to consider gene-mediated mechanisms. Additionally, diseases without treatment do not exist in the largest connected component when genes are removed, so many rare diseases cannot even be embedded.

We expect that the element-wise product of drug and disease vectors representing for known indications should exist closer to the “Treatment” theme in the embedding space than those of randomly-sampled negative indications. To confirm this, we project the combined vectors for pairs in our test set into two dimensions using UMAP.^22^ Figure 3 shows known indications (positive, red) and randomly sampled (negative, blue) drug-disease pairs, where each point is the element-wise product of the drug vector with the disease vector. The positive drug-disease pairs are indeed closer to the “Treatment” vector (black), confirming that our embedding method was able to successfully learn embeddings that reflect a treatment relationship.

**Fig. 3:**
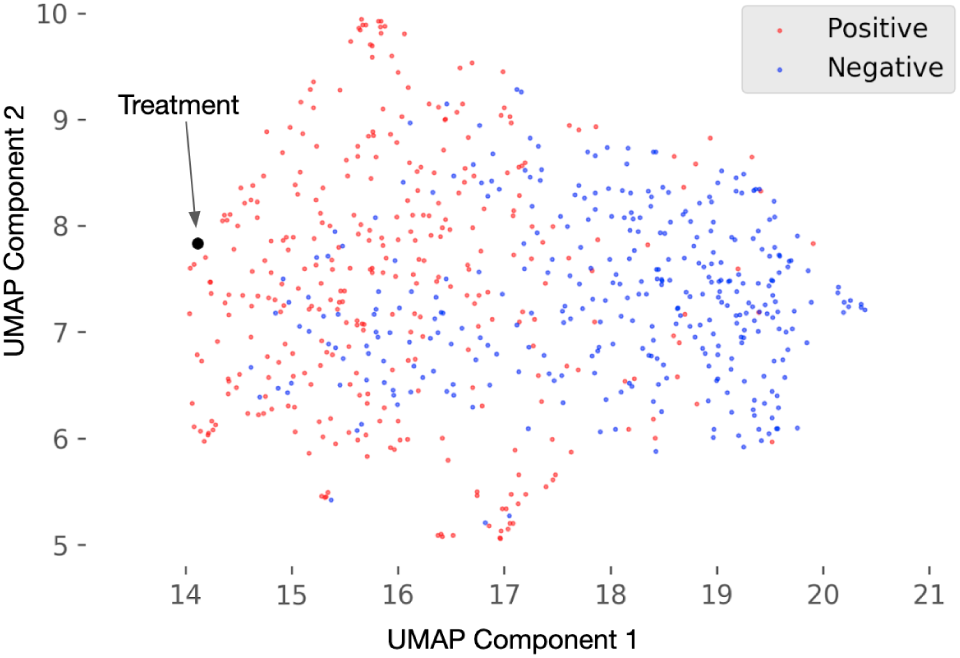
2D UMAP projection of embedded pairs compared to “Treatment”

#### 3.1.3. Evaluation of theme contribution

We hypothesize that the themes which provide the most utility in predicting a treatment-type edge will have embeddings that are similar to that of the “Treatment” theme. To evaluate this, we measured the cosine similarity between the embeddings of all 32 themes in GNBR and perform hierarchical clustering to group the themes. As shown in Fig. 4(a), the theme clusters correspond well with edge type, as expected because the theme embeddings within each group are learned by triples consisting of different node types. Of the two themes which do not appear to cluster with others of the same type, one (E+) belongs to both groups drug (Dr)–gene (G) and G-G. The sub-clusters within each edge type represent biologically relevant internal structure, giving us confidence in the quality of our embeddings. For example, blocks consisting of {“causal mutations” (U), “polymorphisms alter risk” (Y), “mutations affecting disease course” (Ud)} and {“decreased expression” (E-), “antagonism” (A-), “inhibition” (N)} represent important classes of gene-disease and drug-gene relationships, respectively.

**Fig. 4:**
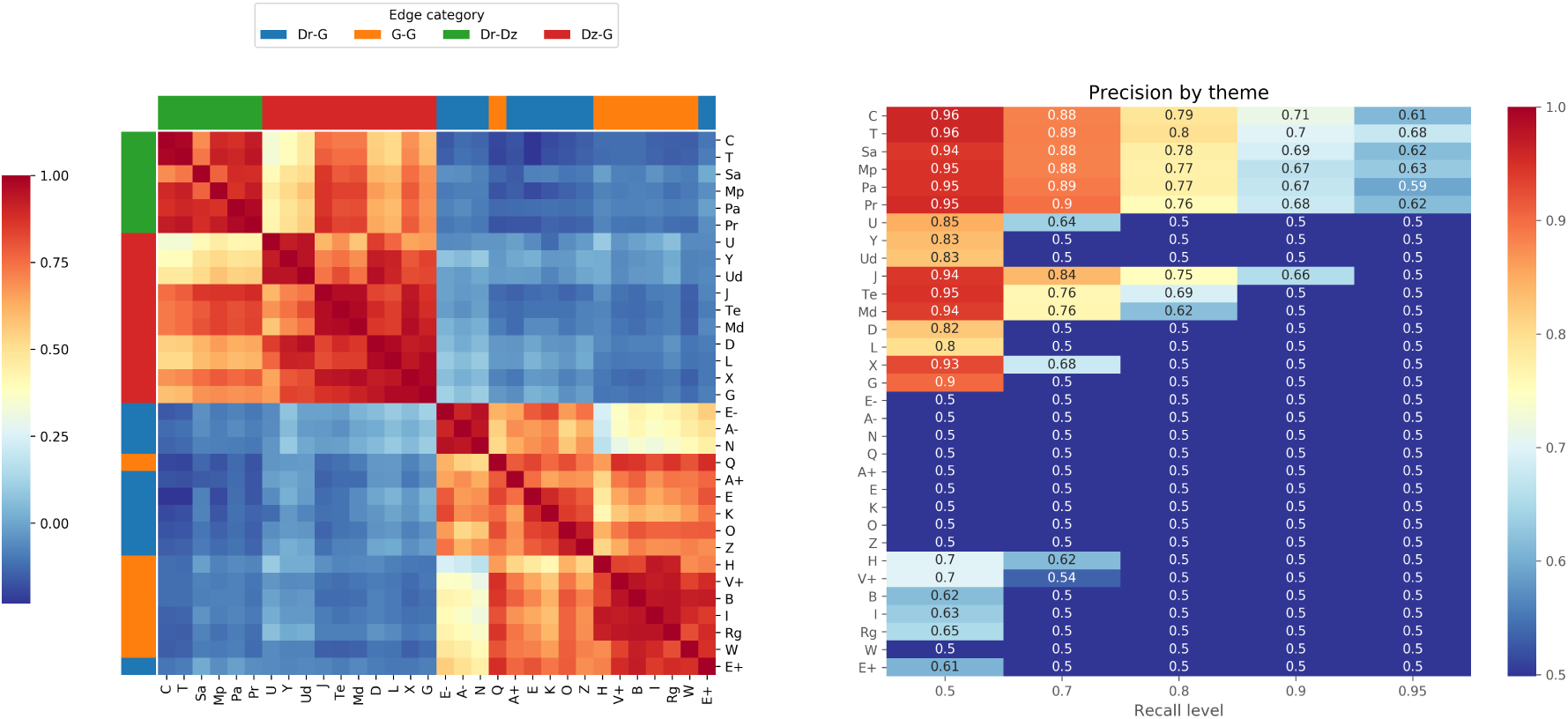
(a) Pairwise cosine similarity (ranging from 1.00 to −0.23) between embeddings for each theme, clustered by hierarchical clustering. (b) Precision at various recall levels for drug-disease score predicted by each theme.

The top-left corner of Fig. 4(a), the Dr–disease (Dz) themes, is particularly relevant for drug repurposing. Note that the similarity between all themes in this group is high, especially “Treatment” (T) and “Inhibits cell growth (esp. cancer)” (C), because of the high literature bias towards cancer compared to other diseases. “Side effect” (Sa) edges are slightly more distant from T and C, which is promising because a side-effect phenotype should not be misconstrued as a treatment indication. In general, the high similarity among all themes in this group suggest that correlation between Dr-Dz themes is indeed the reason for the drop in performance when non-”Treatment” edges are removed from the embedding model. Outside of the Dr-Dz group, we see that among Dz-G themes, three stand out as quite similar to the Dr-Dz themes, suggesting a higher utility in our drug repurposing model (excluding J, which also belongs to the Dr-Dz group): “possible therapeutic effect” (Te), “diagnostic biomarkers” (Md), and “overexpression in disease” (X). Surprisingly, “Drug targets” (D) appears to be less related to treatment, suggesting that the literature references in this category are less specific (e.g. stating that a gene can be targeted without mentioning any actual drug).

To more directly assess how different themes contribute to the prediction of treatment-type edges, we measured the confidence *f* (*l*) for all drug-disease pairs relative to each theme using Eq. (3). The resulting predictions for each theme refer to the likelihood that an edge represents a relationship of that type, but by comparing these predictions to the true labels (i.e. known treatment indications), we can assess the degree to which each contributes to the prediction of a treatment-type edge. Figure 4(b) shows the precision at various recall levels for predicting known treatments. We choose precision as our primary performance metric because for drug repurposing we are most concerned with minimizing false positive predictions. As expected, the “Treatment” theme has the best performance on this task, especially at high recall levels, while other Dr-Dz themes are also highly predictive. The segregation between Dr-Dz/Dz-G and Dr-G/G-G edges suggests that disease nodes are the main drivers of the embeddings and that disease-related edges contain the majority of information relevant to treatment. In accordance with the embedding similarity results, we find that Te, Md, and X are the most predictive non–Dr-Dz themes for disease treatment. These themes may capture recurring semantic patterns that suggest treatment even when such a relationship is not directly stated.

### 3.2. Inferring novel treatments for high-priority rare diseases

The 30 highest-scoring novel drug repurposing candidates for high-priority rare diseases are shown in Table 1. We performed a detailed survey of literature evidence and assess the validity of the prediction using six categories: (1) published treatment, where there is literature evidence indicating the use of the drug to treat human subjects; (2) symptom management, where the drug has been used to address symptoms of the disease; (3) co-morbidity treatment, where the drug treats a comorbid or closely related disease; (4) potentially feasible treatment, where there lies pre-clinical and/or biologically tractable evidence for the drug targeting the rare disease; (5) possible contraindication, where the drug may produce a physiological effect opposite that which is desired; and (6) unknown/no effect, where there is little or no literature evidence to support the drug-disease combination.

**Table 1:**
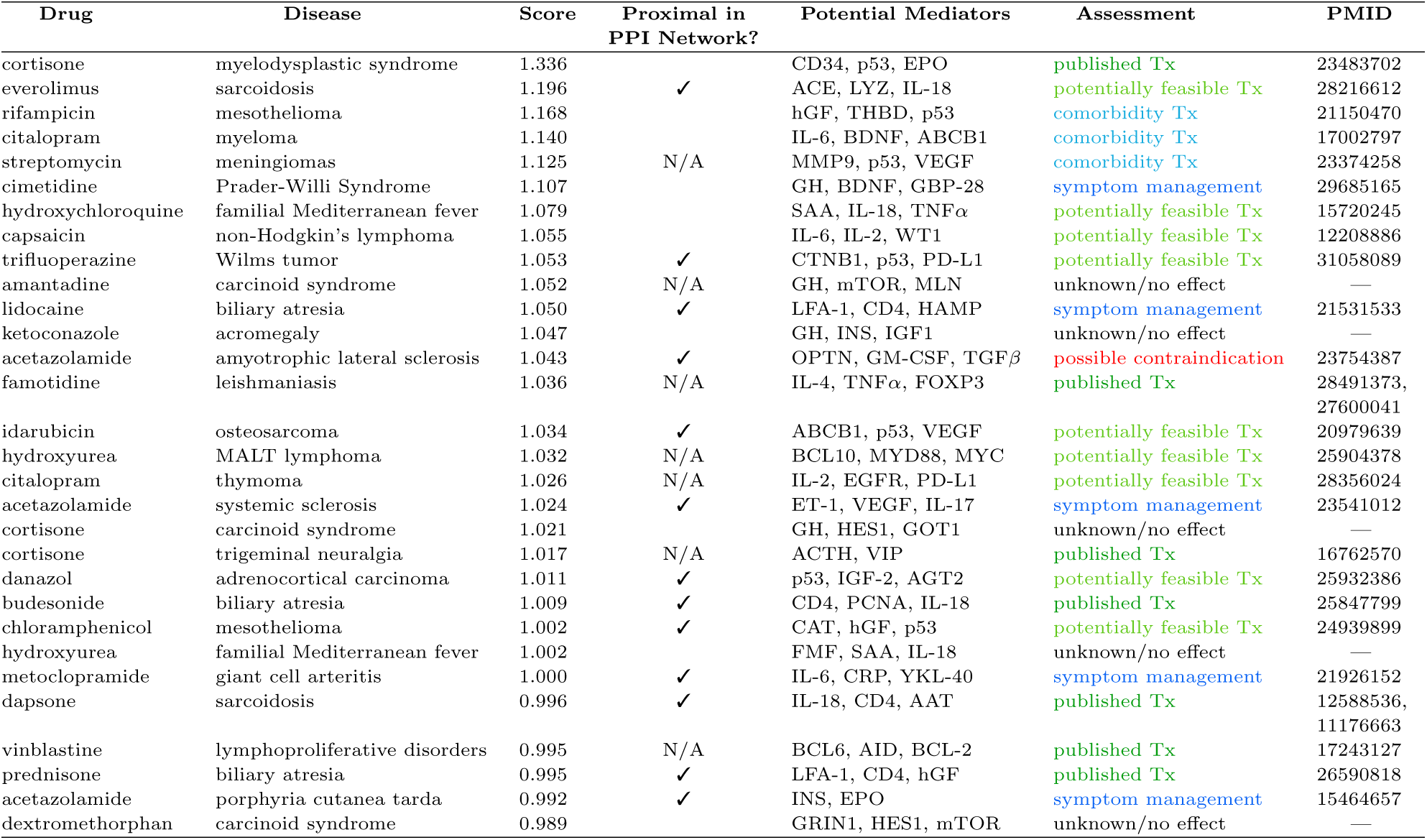
Summary of the top 30 drug repurposing candidates. “Score”: the predicted confidence generated by our model; “Proximal in PPI Network?”: indication of significant proximity between drug- and disease-associated genes (Section 3.3); “Potential Mediators”: top three genes implicated in path analysis (Section 3.4); “Assessment”: manual designation of treatment (Tx) viability; “PMID”: literature reference supporting interpretation.

Most evidence for the feasibility of drug repurposing candidates is not found in a papers abstract or title (the inputs to our model), but within the full text itself. The examples in category 1 provide evidence that our method is able to correctly identify treatments that have been published but are not present in GNBR. The remaining categories represent more complicated cases and demonstrate the additional inductive capacity of our model. Symptom management is one such category; while these drugs do not treat the underlying condition, they can be effective in rare disease patients. For instance, cimetidine, an H2 antagonist that reduces acid in the stomach, may alleviate the gastric reflux symptoms that accompany Prader-Willi syndrome. Other common forms of symptom management drugs include pain relievers, diuretics, and anti-arrhythmic drugs.

Likewise, common comorbidities of rare diseases are not direct treatments, but these cases demonstrate that the model is able to learn patterns in disease co-occurrence and identify drugs that may have a secondary benefit for the rare disease. In some cases, treating the comorbidity may even prevent the onset of the rare disease in question. For example, it is known that tuberculosis (TB) is associated with an increased risk for cancers of the respiratory system such as mesothelioma. As such, drugs that treat TB such as rifampicin may indirectly be protective against mesothelioma.

Possible contraindications represent cases in which our model fails to accurately recognize the nature of the relationship between biological entities. The only possible contraindication noted in the top 30 was between acetazolamide (ACZ) and amyotrophic lateral sclerosis (ALS). To understand why this potential mistake was made, we enumerated all paths of four or fewer nodes containing ACZ and ALS in the largest connected component GNBR graph. Each edge in these paths was reduced to its highest-scoring theme, and the minimum across these theme support scores was used to rank the paths. The highest-ranking path by this method was the following: (ACZ) – [T (0.937)] – (Glaucoma) – [U (0.904)] – (OPTN) – [U (0.906)] – (ALS). In other words, ACZ treats glaucoma, which shares a causal mutation in the OPTN gene with ALS. This suggests that the model correctly identified a mechanistic similarity between the two diseases, but in this case, the conclusion that ACZ would treat both was inaccurate since ACZ does not target OPTN. This is an inherent limitation of high-level semantic knowledge graphs, and incorporating more granular phenotypic effects (e.g. Drug A increases blood pressure) and directionality in edges could lead to a greater ability to learn true mechanistic hypotheses.

Our final category, which we denote as “potentially feasible treatment”, consists of drugs which have been used in pre-clinical or animal studies of the disease, or those that target biological mechanisms influential to the cause or progression of the disease. These examples could lead to novel discoveries and thus represent the most promising candidates for further clinical study. We explore two such cases in more detail.

#### 3.2.1. Case Study 1: Trifluoperazine as a treatment of Wilms tumor

Wilms tumor is a childhood cancer of the kidney. Mutations of the WT1 gene (Wilms Tumor transcription factor gene 1) are responsible for about 20% of Wilms tumor cases.^23^ In particular, WT1 is believed to regulator proto-oncogenes such as MYC in renal development.^24^ Thus, aberrant expression of WT1 can precipitate MYC-mediated cancers. Trifluoperazine is traditionally an antipsychotic but has recently been shown to have anticancer growth properties.^25^ In particular, it is believed to inhibit MYC-induced cell transformation.^26^ We hypothesize that trifluoperazine’s anti-cancer properties can therefore be used to treat cancers in which MYC is dysregulated, such as Wilms tumors. The ability of our method to capture this off-label indication is promising and suggests that the model is learning information from genes proximal to the drug and disease when predicting treatment relationships.

#### 3.2.2. Case Study 2: mTOR inhibition as a treatment of sarcoidosis

Sarcoidosis is a multi-system autoimmune disease with unknown etiology that leads to clusters of inflammatory cells called granulomas in several organs including lungs, skin, and lymph nodes.^27^ mTORC1 pathways activation is a hallmark of these clinical findings. In fact, inhibiting mTOR via drugs, such as everolimus and rapamycin, has slowed down granuloma formation in preclinical animal studies.^28^ To our knowledge, no papers have specifically referenced a relationship between everolimus to sarcoidosis; nonetheless, our method was able to generate a treatment hypothesis based on previous treatment mechanism (mTOR inhibition) and the pathogenesis of the disease (granuloma formation). This suggests that the model is able to synthesize the heterogeneous types of information in the knowledge graph in a way that is meaningful for evaluating treatment potential.

### 3.3. External validation using network proximity

To externally validate the pairs in Table 1, we calculated the network proximity between sets of drug target genes derived from DGIdb^29^ and disease-associated genes derived from OrphaNet and OMIM. The underlying assumption is that gene sets corresponding to true drug-disease combinations will be closer to each other in a protein-protein interaction (PPI) network than expected by random chance. We represented every node in the STRINGdb v10^30^ PPI network as a 128-dimensional embedding vector using Node2Vec^31^ with default parameters and calculated the median cosine similarity between proteins in the drug and disease sets.^32^ We calculated an empirical *p*-value based on 10,000 samples of genes drawn randomly with replacement into two sets of the same size as the true drug and disease sets. Seven pairs from Table 1 could not be calculated because either the drug or disease did not have a corresponding gene set. Thirteen of the remaining 23 pairs were found to be significant under a Bonferroni-adjusted *p*-value threshold of 0.05, demonstrating that many of our predictions can be corroborated by independent data under this network proximity hypothesis. Those that do not pass the significance threshold may simply not be related by a known genetic mechanism; failure to identify network proximity does not preclude a true treatment, especially for entirely novel predictions.

### 3.4. Drug-disease path analysis

To better understand how our model makes novel link predictions, we analyzed the four-node paths connecting drug-disease pairs in the original GNBR graph, ranked using the procedure described in the ACZ-ALS example above. There are three possible metapath motifs based on node type: drug-disease-gene-disease (DzG-mediated), drug-disease-drug-disease (DzDr-mediated), and drug-gene-gene-disease (GG-mediated). In the first, the drug treats a different disease with the same genetic mechanism; in the second, the drug treats a different disease which shares a treatment with the disease of interest; and in the third, the drug affects the disease via two interacting genes. A representative example of each is shown in Fig. 5(a).

**Fig. 5:**
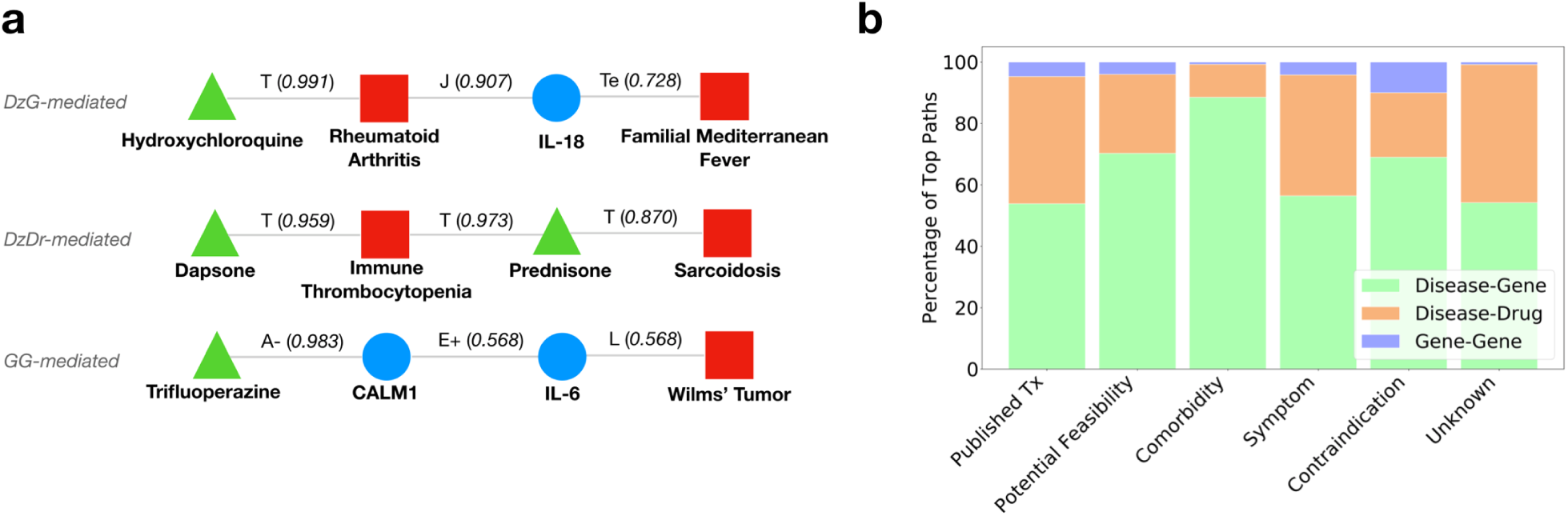
(a) Examples of each drug-disease path motif. Edges are labeled with their highest-supported themes and corresponding support scores. (b) Distribution of motifs across the six interpretation categories in Table 1 as determined by the occurrence of each motif across the top 100 ranked paths per drug-disease prediction.

In particular, GG-mediated paths demonstrate the models ability to automatically identify biological mechanisms that do not rely transitively on other diseases. In the example shown in Fig. 5, we show an alternative, immunological hypothesis for the relationship in Case Study 1 (section 3.2.1). Here, trifluoroperazine is known to antagonize calmodulin (CALM1), a calcium-binding protein that induces expression of the inflammatory cytokine interleukin-6 (IL-6). Improper regulation of IL-6 is implicated in the progression of Wilms tumor,^33^ suggesting that antagonizing calmodulin could indirectly help slow tumor progression by reducing IL-6 expression. Cases such as this exemplify the capacity of our model to not only identify new treatment opportunities, but also develop a more nuanced understanding of how these treatments may manifest if they are successful.

Finally, we analyzed the distribution of path motifs for each of our defined feasibility categories to discover any systematic patterns in how the model infers link predictions (Fig. 5(b)). We observe a relative enrichment in the DzG-mediated motif for pairs classified as comorbidity treatment. This is because two diseases that share a common genetic mechanism are frequently comorbid, and the model predicts that a drug that treats one of the conditions also treats the other. This can be a useful inference, especially in cases where the drug affects the mediating gene, but if not it can result in mistakes like the ACZ-ALS case. Such errors could be mitigated by incorporating information about the drugs mechanism of action (e.g. drug class) during inference to rule out obvious mismatches. Published treatments had fewer gene-mediated connecting paths because the evidence for these relationships is mostly contained in clinical journals, which do not typically discuss mechanism in the abstract. GG-mediated motifs were also infrequent in the top-ranked paths, most likely because a drug’s target genes are not referenced in abstracts as much as its disease indications, and thus drug-gene edges tend to have lower scores in GNBR. This suggests that a GNBR-like knowledge graph derived from full texts as well as abstracts could provide even more power to discover novel relationships.

More generally, the capacity of our approach to learning new treatment is dependent on the information represented in the underlying knowledge graph. In the case of GNBR, we rely on a fixed set of themes extracted from unsupervised clustering in a manner agnostic to the downstream task. Our method may benefit from a more granular, fine-tuned set of themes that better encapsulate important information for drug repurposing. Additionally, as NLP tools improve in their ability to capture complex relationships across unstructured text (e.g. dependencies across sentences), knowledge graphs such as GNBR will become more comprehensive, and our methods capacity to learn patterns across the literature will increase.

## 4. Conclusion

We describe a method for generating drug repurposing hypotheses for these rare diseases using embeddings learned from the GNBR knowledge graph. Our approach is fully automated and takes advantage of the vast amount of unstructured information across the medical literature, while explicitly modeling the confidence in this information. We demonstrate high performance on a gold standard set of drug indications, as well as the ability to generate novel drug repurposing hypotheses. We further provide evidence to support our treatment predictions using independent sources, and identify specific motifs in the original knowledge graph which help explain model behavior. Our model was able to successfully learn biologically-relevant patterns from noisy knowledge-based data, but in the absence of experimental validation these predictions remain simply hypotheses. However, our approach of automatically synthesizing literature knowledge helps to narrow the search space of clinical research and thus accelerate the discovery of new treatment options for patients with rare diseases.

## 5. Acknowledgments

This work originally began as a class project for BIOMEDIN 212, “Introduction to Biomedical Informatics Research Methodology”, at Stanford University. We would like to thank the teaching staff, Hunter Boyce and Erika Strandberg, for their constructive feedback, as well as Stefano Rensi for his advice and GNBR technical support. D.S. is supported by LM007033, A.D. is supported by LM012409, and M.G. is supported by GM007365. R.B.A. is supported by NIH LM005652 and GM102365.

